# Differential Pathogenic and Commensal Response of *Staphylococcus aureus* and *Staphylococcus epidermidis* Towards Chemical Signals of Human Skin

**DOI:** 10.64898/2026.01.05.697747

**Authors:** Tika Bahadur Thapa, Robert C. Kuiack, Martin J. McGavin

## Abstract

Pathogenic *Staphylococcus aureus* and commensal *Staphylococcus epidermidis* encounter acidic pH and C16 fatty acids on human skin, but *S. aureu*s uniquely has a complete *fad* pathway for metabolism of saturated C16:0 palmitic acid. We now report on significant differences in their response to C16 fatty acids during growth at pH 5.5. Unsaturated palmitoleic acid C16:1 was more toxic to *S. aureus*, but toxicity was mitigated by saturated C16:0. Consistent with a functional *fad* pathway, C16:0 conferred enhanced growth to *S. aureus*, but not *S. epidermidis*. Acidic pH and C16 fatty acids stimulated SspA serine protease production in *S. aureus* but repressed the orthologous Esp protease in *S. epidermidis*. Although *S. aureus* biofilm formation was stimulated by acidic pH and C16:0, this effect was abrogated by 25 µM C16:1 which promotes protease production, whereas *S. epidermidis* maintained enhanced biofilm in presence of C16:1. Exogenous C16:0 was directly incorporated into phospholipid by *S. epidermidis* but was extended to C18:0 and C20:0 in *S. aureus* prior to incorporation. This may account for differential signaling through the GraSR two component sensor, which is required for SspA production in *S. aureus* at acidic pH. Notably, singular *graSR* dependent phenotypes in *S. aureus graS/R* deletions were restored by *S. epidermidis graS/R* at acidic pH alone, whereas growth at pH 5.5 with 25 µM C16:1 could only be restored with *S. aureus graS/R*. These findings provide important new insight into how members of the Staphylococcal genus are differentially influenced by common environmental signals on human skin.

**IMPORTANCE:** Human skin is a chemically hostile environment, with acidic pH and antimicrobial fatty acids that challenge microbial survival. Understanding how closely related *Staphylococcus epidermidis* and *Staphylococcus aureus* navigate these conditions is critical for distinguishing commensal behavior from pathogenic potential. Our research reveals that *S. aureus* and *S. epidermidis*, though genetically similar, employ markedly distinct adaptive mechanisms in response to identical skin-derived cues. Specifically, each species remodels its membrane phospholipids in unique ways under acidic pH and C16 fatty acid exposure. These environmental factors also differentially modulate their biofilm formation and protease activity. Together, our findings highlight how the same host-derived chemical signals of skin can activate virulence-associated traits in *S. aureus* while supporting commensal persistence in *S. epidermidis*.

## INTRODUCTION

For members of the *Staphylococcus* genus that colonize human hosts, the skin constitutes a unique ecological niche defined by its mildly acidic pH, limited nutrients, and free fatty acids (FFAs). Under healthy conditions, skin surface pH typically ranges from 4.1 to 5.8, a barrier that both restricts pathogen colonization and shapes commensal communities (1, 2). Concurrently, sebaceous glands secrete significant amounts of C16 FFAs, of which unsaturated sapienic acid (C16:1n-6) and palmitoleic acid (C16:1n-9) exert broad antimicrobial effects via membrane disruption and enzyme inhibition (3, 4), whereas saturated palmitic acid (C16:0) represents a potential nutrient that could be utilized through the recently described *fad* pathway for β-oxidation in *Staphylococcus aureus* (5), which is incomplete in *Staphylococcus epidermidis*. Nevertheless, *S. epidermidis* is a ubiquitous commensal of human skin, whereas *S. aureus* primary site of colonization is the anterior nares of approximately 25-30% of humans, which is requisite to subsequent colonization of skin (6–8). While the differential ability of *S. epidermidis* and *S. aureus* to colonize human skin is well documented, differences in their ability to sense and respond to the associated common environmental cues is less well defined, especially in context of their being closely related species that share many orthologous cell surface and secreted proteins, and their associated regulators.

As a predominant constituent of the healthy skin microbiome, *S. epidermidis* contributes to barrier integrity and immune modulation through specialized metabolites and enzymatic activities (9, 10), whereas *S. aureus* is an opportunistic pathogen whose presence on skin is often associated with disease, particularly in atopic dermatitis and wound infections (11, 12). *S. aureus* virulence is orchestrated by intricate regulatory networks including the accessory gene regulator quorum sensing system (*agr*), and two-component systems such as GraRS, SaeRS and ArlRS, (13–15), all of which are also maintained in *S. epidermidis* (16–19). In *S. aureus*, we discovered that signaling through the GraS sensor kinase was activated by acidic pH, under which condition it was required for expression of the SspA serine protease and conferred enhanced resistance to linoleic acid, adding to its previously documented activation by antimicrobial peptides (20). Namely, signaling through GraS activates the GraR response regulator, leading to expression of genes including *mprF*, and *dltABCD* which mediate resistance to cationic peptides by modifying membrane charge and composition (21, 22). However, orthologous genes are also found in *S. epidermidis*, where GraRS exhibits similar structural organization and function in sensing and responding to cell envelope stress (17).

Another shared pathway relevant to host interaction is inherent in utilization of host-derived free fatty acids to support phospholipid synthesis through the fatty acid kinase FakAB pathway (5, 23, 24). FakA phosphorylates FFAs bound by FakB proteins, facilitating either direct acyl chain incorporation into phospholipid through the PlsY acyl transferase, or alternately being submitted to the type II fatty acid biosynthesis (FASII) elongation cycle prior to incorporation into phospholipid through the combined actions of PlsX which catalyzes the transfer of fatty acids between phosphate and acyl carrier protein (ACP), and the PlsC acyl transferase that utilizes acyl-ACP substrates (25). Membrane remodeling through incorporating these fatty acids has profound biophysical consequences, as saturated fatty acids promote tight acyl chain packing and increased membrane rigidity, while even minor incorporation of monounsaturated chains like palmitoleate introduces kinks that disrupt lipid order and enhance fluidity (26–28). Therefore, while it is feasible that even minor differences in signaling through orthologous regulators, or functions of shared metabolic processes may promote different outcomes in support of commensal or pathogenic phenotypes, this avenue of investigation has remained largely undeveloped.

An intriguing example relates to the function and regulation of orthologous serine proteases SspA (V8 protease) of *S. aureus* and Esp of *S. epidermidis*, which are both members of the glutamyl endopeptidase family. SspA is a virulence factor in atopic dermatitis, promoting itchy skin through proteolytic activation of the PAR1 receptor (29), and consistent with this role in virulence, our work has revealed that SspA expression is stimulated by acidic pH and unsaturated fatty acids that would be encountered on human skin in a GraS-dependent manner (20). Conversely, the orthologous Esp of *S. epidermidis* is proposed to interfere with *S. aureus* colonization of humans through disruption of biofilm formation (30). Herein, we have assessed protease production in *S. aureus* and *S. epidermidis* as one marker for how these bacteria may differentially respond to environmental signals encountered on human skin, with emphasis on acidic pH and C16 fatty acids. Our approach integrated growth kinetics, phospholipid composition, and heterologous GraRS complementation. Strikingly, whereas heterologous GraRS complementation in *S. aureus* revealed some conservation of function with respect to modification of membrane charge, we nevertheless find that growth, protease production, biofilm formation and phospholipid composition are all differentially modulated in *S. aureus* and *S. epidermidis* in response to environmental cues of acidic pH and C16 fatty acids.

## RESULTS

### Growth of *S. aureus* and *S. epidermidis* is differentially influenced by acidic pH and host derived fatty acids

To persist on skin, bacteria must contend with both acidic pH and antimicrobial unsaturated free fatty acids, of which sapienic acid (C16:1n-6) and its isomer palmitoleic acid (C16:1n-9) are most abundant in sebaceous secretions, while palmitic acid (C16:0) which is normally not toxic to *S. aureus* is the most abundant saturated fatty acid (31, 32). To determine the extent to which *S. aureus* USA300 and *S. epidermidis* M23864:W2 are differentially influenced by these chemical signals, we first assessed growth in glucose-free TSB with modifications, including buffered at pH 5.5, or pH 5.5 with combinations of palmitic (C16:0) and palmitoleic acid (C16:1). Surprisingly, *S. epidermidis* failed to grow over 8h in unbuffered TSB + 500 µM palmitic acid, but this restriction was not evident at pH 5.5 (Fig 1A, B), while growth of *S. aureus* was not restricted in either TSB or TSB pH 5.5. Moreover, consistent with it having a functional *fad* pathway for metabolism of palmitic acid through β-oxidation, palmitic acid promoted a higher 24h growth yield for *S. aureus* in both unbuffered and pH 5.5 media, whereas no such increase was evident in *S. epidermidis* (Fig S1A, B), which has an incomplete *fad* pathway.

**Fig 1:**
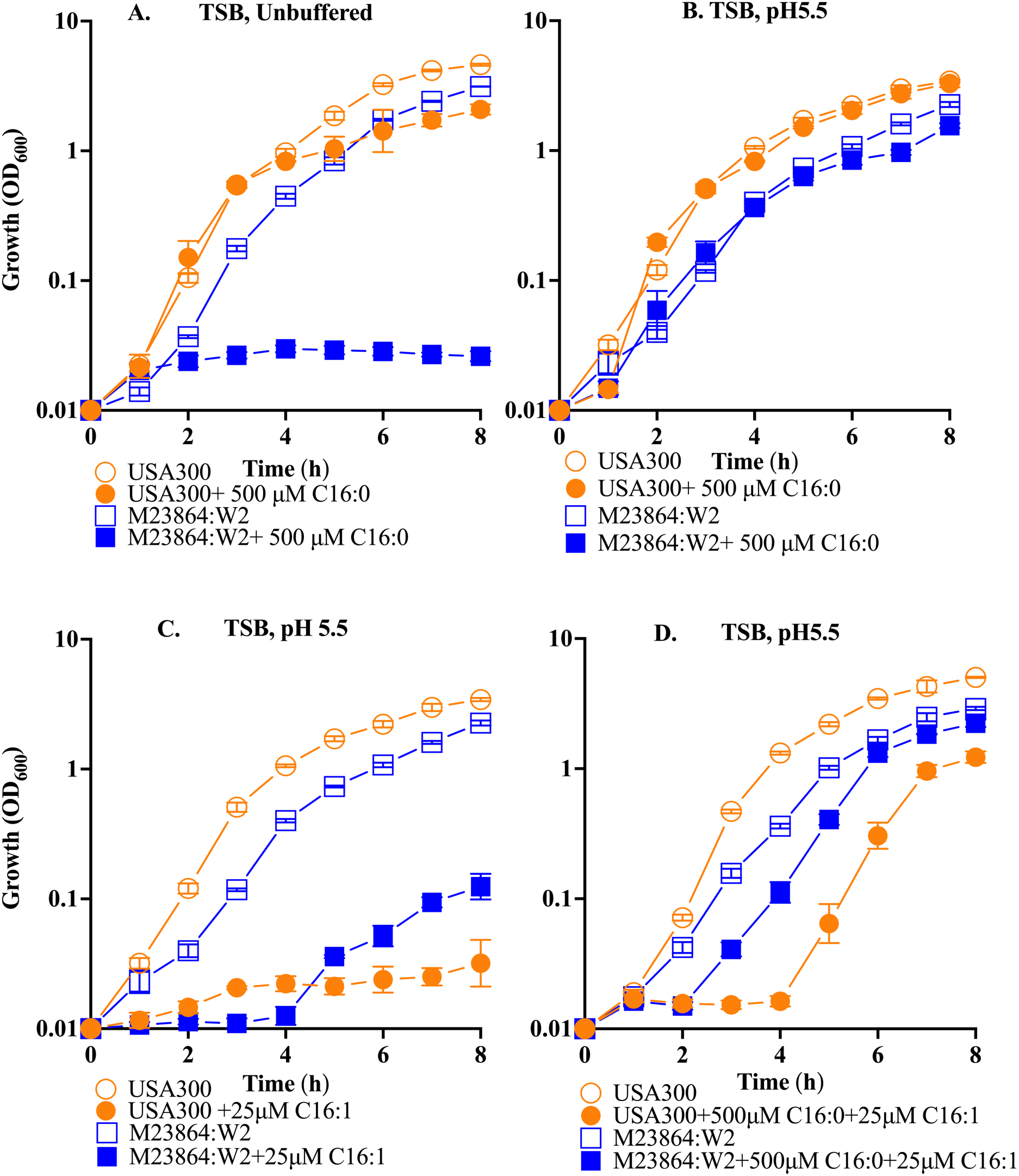
Influence of acidic pH and C16 fatty acids on growth of *S. aureus* USA300 and *S. epidermidis* M23864:W2. Triplicate flasks of TSB (unbuffered A, or buffered at pH 5.5 B, C, D) were supplemented with indicated concentrations of palmitic acid C16:0 (A, B), palmitoleic acid C16:1 (C), or C16:0 + C16:1(D) and inoculated to an initial OD_600_ of 0.01. Growth (OD_600_) was assessed at hourly intervals, and each data point represents the mean±SEM from triplicate flasks.

Reflecting the toxicity of unsaturated fatty acids, *S. epidermidis* exhibited a 4h lag phase in TSB pH 5.5 + 25 µM palmitoleic acid, and *S. aureus* exhibited a prolonged lag phase (Fig 1C). However, the toxicity of 25 µM palmitoleic acid towards both microbes at pH 5.5 was mitigated by inclusion of 500 µM palmitic acid, under which condition *S. epidermidis* initiated growth after a 2h lag, while *S. aureus* exhibited a 4 lag and a greater disparity in growth compared to TSB pH 5.5 alone relative to *S. epidermidis* (Fig 1D). At 24h, *S. aureus* and *S. epidermidis* exhibited comparable growth in TSB pH 5.5 + 25 µM palmitoleic acid, whereas in TSB pH 5.5 with both palmitoleic and palmitic acid, growth of *S. aureus* was once again enhanced relative to *S. epidermidis* (Fig S1C, D).

We also assessed the influence of acidic pH on minimum inhibitory concentration (MIC) of palmitoleic acid, which was 100 µM and 200 µM for *S. aureus* and *S. epidermidis* respectively in unbuffered TSB, increasing to 400 µM and > 1600 µM in TSB pH 5.5 (Table S1). Cumulatively, these data reveal differential abilities of *S. aureus* and *S. epidermidis* to grow at acidic pH in the presence of palmitic and palmitoleic acid that would be encountered on human skin. Consistent with its commensal nature, *S. epidermidis* is better adapted to growth at acidic pH in presence of unsaturated palmitoleic acid. However, acid pH and the presence of palmitic acid mitigated the toxicity of palmitoleic acid for both microbes, and with a complete *fad* pathway *S. aureus* is uniquely capable of achieving higher growth yields in the presence of palmitic acid.

### *S. aureus* and *S. epidermidis* exhibit distinct differences in phospholipid composition in response to acidic pH and exogenous fatty acids

Although *S. aureus* and *S. epidermidis* have orthologous *fakAB* and other conserved genes for incorporation of exogenous fatty acids into phospholipid, the mechanistic details have been primarily elucidated in *S. aureus* where endogenously synthesized fatty acids exit the FASII pathway on an acyl carrier protein ACP, after which PlsX catalyzes the conversion of acyl-ACP to acyl-phosphate, which is then used by acyl transferase PlsY to acylate the sn-1 position of glycerol-3-phosphate, producing 1-acyl-G3P. Another acyl transferase PlsC directly utilizes acyl-ACP from the FASII pathway, with a strong preference for branched chain C15 fatty acid, to acylate the sn2 position of 1-acyl-G3P, forming phosphatidic acid, the metabolic precursor of phosphatidylglycerol (33, 34). Exogenous fatty acids that enter the cytoplasm are bound by FakB and phosphorylated by FakA. The acyl-phosphate is then used by PlsY to acylate the sn1 position of G3P as described for the endogenous pathway (35). However, in *S. aureus* PlsY does not function effectively with C16, such that acyl(C16)-phosphate is converted to acyl-ACP *via* PlsX and submitted to FASII for extension to C18 and C20, after which PlsX converts the extended acyl-ACP to acyl-phosphate, which is then used by PlsY to acylate the sn-1 position of G3P. Therefore, when *S. aureus* is grown with exogenous C16 fatty acid, it is extended to C18 or C20 via FASII prior to incorporation into the sn1 position of G3P, and the stringent preference for incorporation of endogenously synthesized branched chain C15 at the sn2 position is maintained (23, 24, 33). However, it is not known if these specificities are maintained at acidic pH, and whether *S. epidermidis* is subject to these same restrictions.

In previous studies with *S. aureus* grown in TSB, the PG species predominantly contained branched chain fatty acids with PG32:0 as the major species, comprised of C15 at sn2 and C17 at sn1 (5, 24). Comparatively, when we assessed phospholipid composition from cells grown in TSB pH 5.5 with no exogenous fatty acid, PG32:0 in both *S. aureus* and *S. epidermidis* comprised just 10% of the total PG (Fig 2), while PG33:0 (C15 + C18) was a major component for both bacteria at ∼ 37%. However, the composition of other major PG species differed significantly. In *S. aureus* the 37% PG33:0 was accompanied by 32% PG31:0 (C15 + C16) and 18% PG35:0 (C15 + C20), while in *S. epidermidis* the abundances of PG31:0 and PG35:0 were reversed, with just 7% PG31:0 and 43% PG35:0. Consequently, at acidic pH *S. epidermidis* has a greater tendency to incorporate longer saturated straight chain C18 and C20 fatty acids into PG, such that PG33:0 and PG35:0 comprised 79% of total PG, compared to only 55% in *S. aureus*.

**Fig 2:**
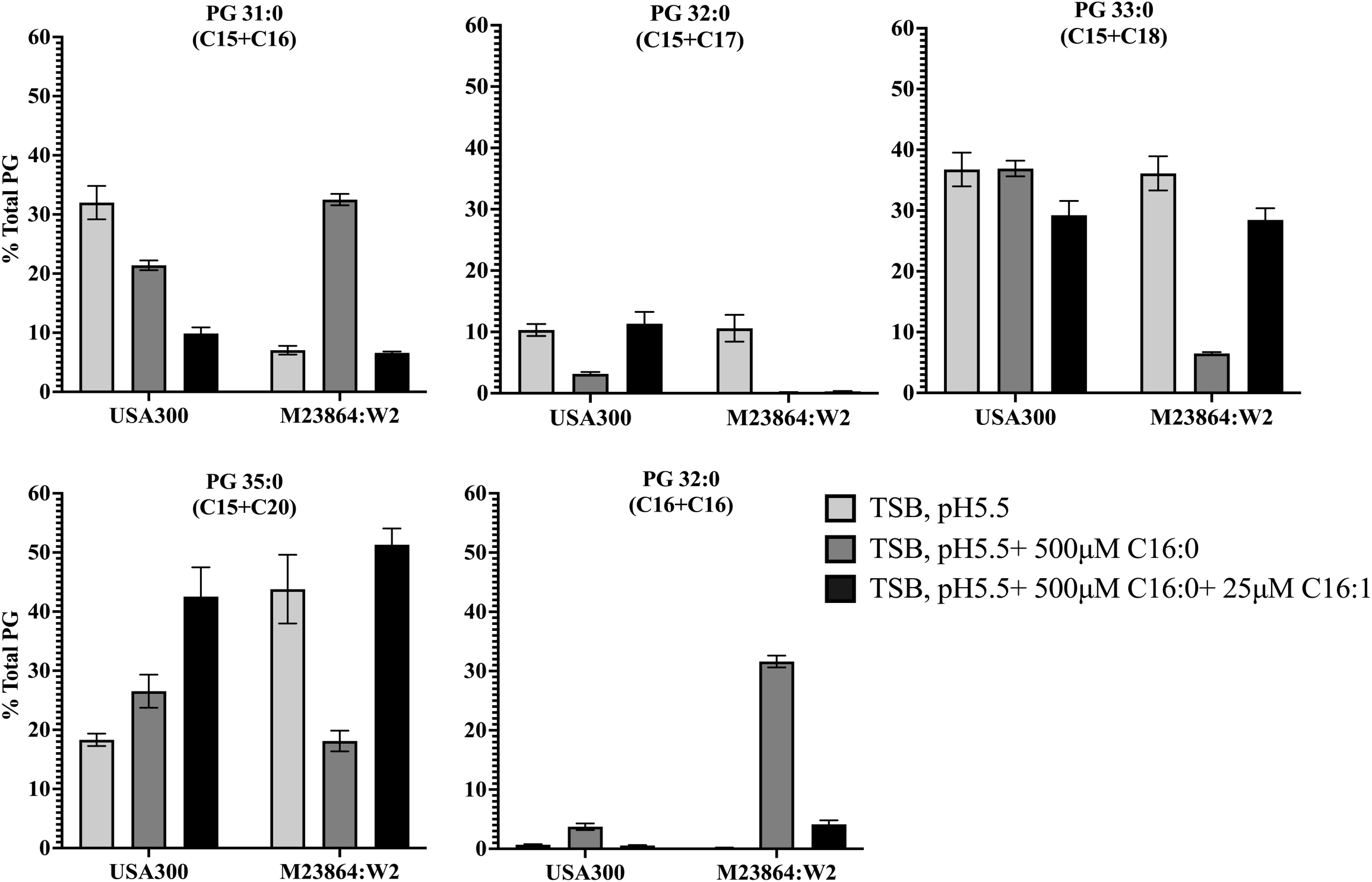
Membrane phospholipid composition of *S. aureus* USA300 and *S. epidermidis* M23864:W2 is differentially affected by acidic pH and exogenous C16 fatty acids. USA300 and M23864:W2 were grown in triplicate cultures to an OD_600_ of 0.6 in TSB pH5.5 alone, TSB pH 5.5 + 500 µM palmitic acid C16:0, or pH 5.5 with both 500 µM palmitic acid and 25 µM palmitoleic acid C16:1 as indicated. Cultures were then processed for extraction and analysis of phosphatidylglycerol (PG) content. Results are presented as mean ± SEM of the percentage of total PG each lipid species represents. PG composition is presented as the sum of the saturated carbon chain lengths at the sn1 and sn2 positions of each PG species as indicated above each graph. Data for minor PG components with C16:1 are depicted separately (Fig. S2).

At pH 5.5 with 500 µM palmitic acid, *S. aureus* exhibited modest changes in PG composition. Strikingly, the amount of PG33:0 was not significantly altered compared to growth at pH 5.5 alone, while a decrease in the amount of PG31:0 was accompanied by an equivalent increase in PG35:0. This is consistent with incorporation of C15 at sn2 being maintained while the exogenous C16 is extended to C20 via FASII prior to incorporation at sn1, accounting for the increase in PG35:0. Surprisingly, whereas the PG33:0 content of *S. aureus* was not altered in response to exogenous palmitic acid, it dropped in *S. epidermidis* from ∼ 37% to 6%, which also displayed a major reduction in PG35:0 from 43% to 18%. These reductions were accompanied by compensatory increases in PG that contained C16 fatty acids, such that PG31 (C15 + C16) and PG32 (C16 + C16) together accounted for 63% of the total PG species. Therefore, in contrast to *S. aureus*, *S. epidermidis* can directly incorporate exogenous C16 fatty acids into phospholipid and is also less restricted in incorporation of C15 at the sn2 position of PG, as evident from PG32:0 (C16 + C16) comprising >30% of the total PG species.

Surprisingly, when both palmitic and palmitoleic acids were supplemented together, the PG profiles of *S. aureus* and *S. epidermidis* converged. In this condition, PG33:0 and PG35:0 comprised approximately 71% and 79% of total PG in *S. aureus* and *S. epidermidis* respectively (Fig. 2). Since palmitoleic acid was only included at 25 µM, its utilization was not sufficient to appear in a major PG species. Nevertheless, *S. aureus* and *S. epidermidis* both displayed some ability to incorporate C16:1 into PG. Notably, *S. aureus* had 1.5% PG31:1 (C15 + C16:1) which was not evident in *S. epidermidis*, whereas *S. epidermidis* had 4% PG36:1 (C20 + C16:1) compared to less than 0.5% in *S. aureus* (Fig S2). These findings highlight species-specific differences in fatty acid incorporation and membrane remodeling in response to acidic pH and C16 fatty acids.

### *S. aureus* and *S. epidermidis* exhibit similar capacity to modulate membrane fluidity in response to acidic pH and exogenous C16 fatty acids

To assess how host-derived fatty acids influence bacterial membrane biophysical properties, we measured membrane fluidity using Laurdan generalized polarization (GP), a dye-based assay sensitive to lipid packing. Higher GP values reflect increased membrane order (rigidity), while lower values indicate increased fluidity. In TSB pH 5.5, *S. aureus* and *S. epidermidis* exhibited similar Laurdan GP values, and when supplemented with exogenous palmitic acid, there was a marked increase in membrane rigidity (Fig 3), consistent with the noted reduction in PG32:0 containing branched chain C15 and C17 fatty acids that promote membrane fluidity, concomitant with increased abundance of PG species containing straight chain C16 fatty acid in *S. epidermidis*, or C20 in *S. aureus*. (Fig 2). These saturated straight chain fatty acids promote tighter acyl chain packing, thus increasing membrane rigidity (36, 37). Co-supplementation of C16:0 with C16:1 reduced GP values in both *S. aureus* and *S. epidermidis* compared to C16:0 alone (Fig 3), though not to the same measure as recorded in the absence of fatty acid. Incorporation of monounsaturated palmitoleic acid into the minor PG species PG31:1 (C15+C16:1) in *S. aureus* and PG36:1 (C20+C16:1) in *S. epidermidis* (Fig S2) likely accounts for reduction in rigidity relative to cultures grown in palmitic acid alone, since even at low abundance, incorporation of unsaturated fatty acids is known to significantly lower membrane order (38). Together, these results demonstrate that *S. aureus* and *S. epidermidis* exhibit a similar capacity to modulate membrane fluidity in response to acidic pH and host derived C16 fatty acids, even though they display significant differences in phospholipid composition. A caveat to this conclusion is that we did not assess higher concentrations of palmitoleic acid, which *S. epidermidis* is better able to tolerate.

**Fig 3:**
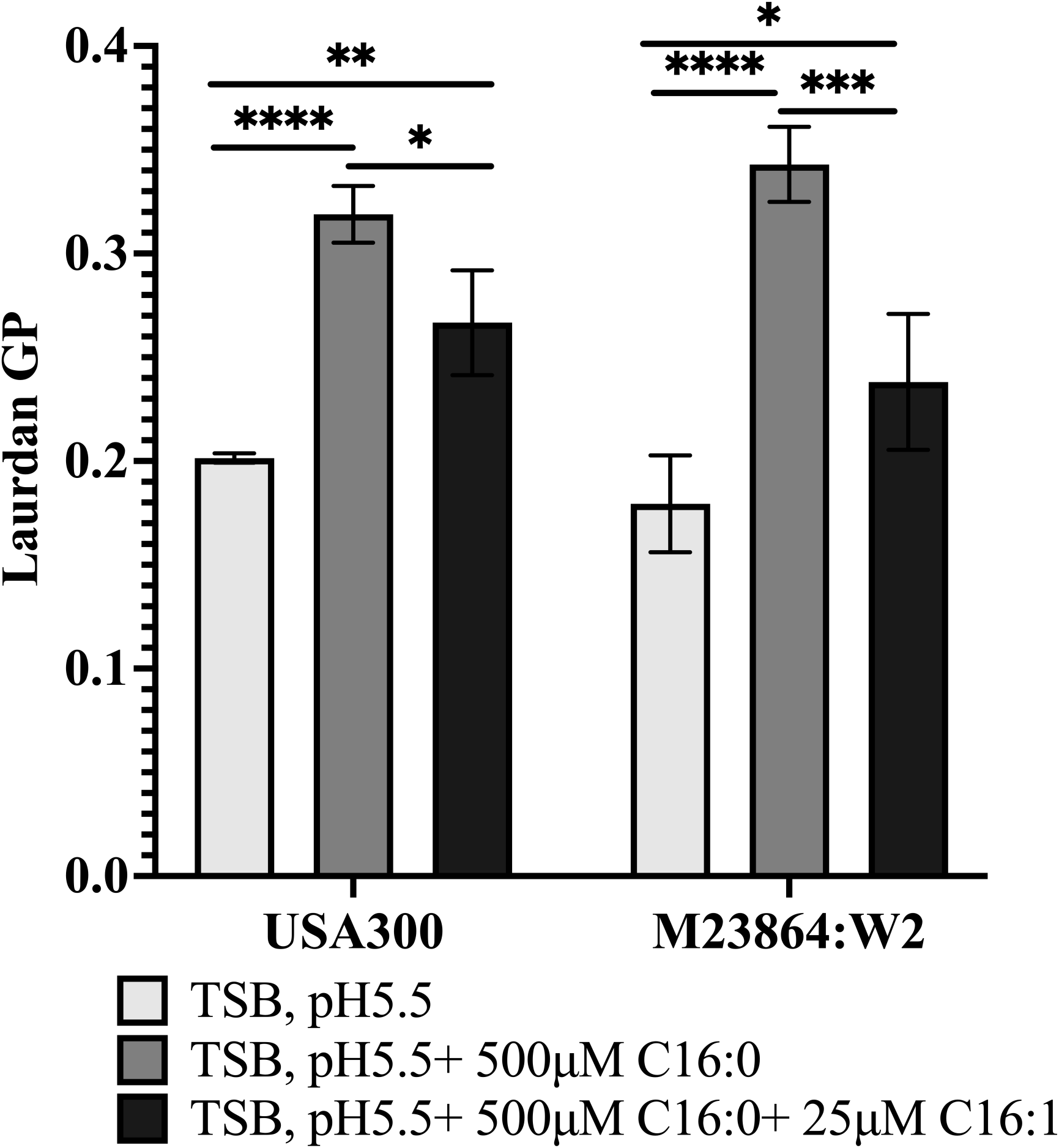
Laurdan generalized polarization (GP) assay to evaluate influence of C16 fatty acids on membrane fluidity in *S. aureus* USA300 and *S. epidermidis* M23864:W2 during growth in TSB pH 5.5. USA300 and M23864:W2 were grown in TSB pH 5.5 alone, or TSB pH 5.5 supplemented with either 500 µM palmitic acid C16:0, or with 500 µM palmitic acid and 25 µM palmitoleic acid C16:1 as indicated, until mid-log phase (OD_600_ = 0.4). Bacterial cells were harvested, stained with laurdan dye, and membrane fluidity was assessed by calculating GP values from fluorescence emission. Higher GP values indicate increased membrane rigidity, while lower values reflect enhanced fluidity. Data represent mean ± SEM from triplicate flasks. Statistical significance was measured using Two-Way ANOVA with Tukey’s multiple comparisons test, *p<0.05, **p < 0.01, ***p < 0.001, ****p<0.0001.

### Host derived fatty acids differentially modulate biofilm formation in *S. aureus* and *S. epidermidis*

Biofilm formation is reported to aid both *S. aureus* and *S. epidermidis* in adhering to epithelial surfaces and withstanding skin’s antimicrobial defenses, including acidic pH and fatty acids (39, 40). We therefore conducted experiments to determine if biofilm formation is differentially influenced by combinations of acidic pH and C16 fatty acids that would be encountered on human skin. Growth of *S. aureus* in TSB pH 5.5 promoted a significant increase in biofilm formation compared to unbuffered TSB, which was abrogated by inclusion of 25 µM palmitoleic acid (Fig 4). In contrast, 500 µM palmitic acid promoted an additional but non-significant increase in biofilm formation compared to pH 5.5 alone, but this stimulatory effect was again abrogated by 25 µM palmitoleic acid. Comparatively, acidic pH alone did not stimulate biofilm accumulation in *S. epidermidis*, whereas acidic pH plus 500 µM palmitic acid promoted a significant increase, which was maintained in presence of 25 µM palmitoleic acid. Therefore, acidic pH combined with palmitic and palmitoleic acid as would be encountered on human skin, stimulates biofilm formation in *S. epidermidis* but not *S. aureus*.

**Fig 4:**
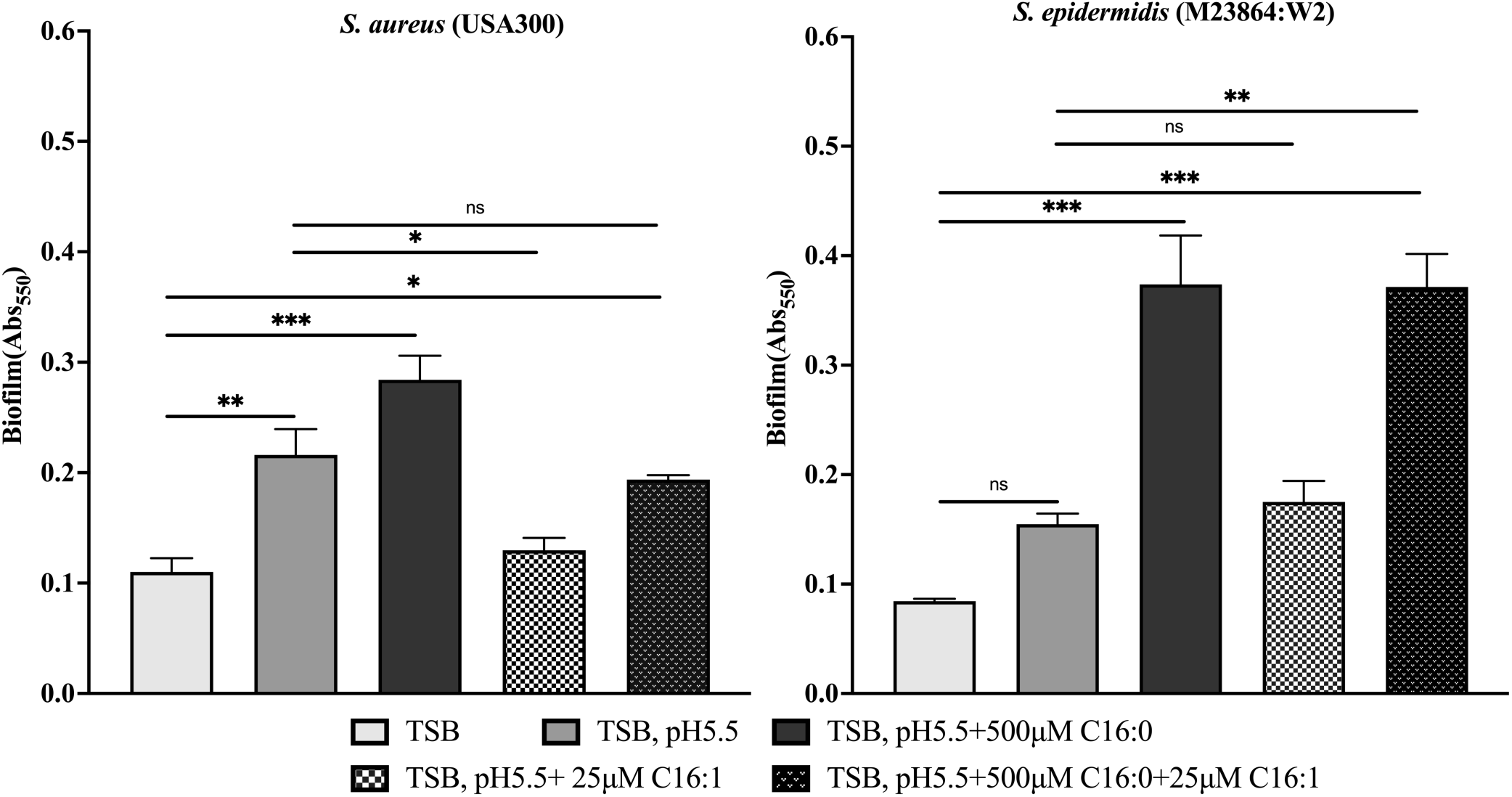
Influence of acidic pH and C16 fatty acids on biofilm formation by *S. aureus* USA300 and *S. epidermidis* M23864:W2. USA300 and M23864:W2 were inoculated to OD_600_ of 0.01 in wells of microtiter plates containing 200 µl of unbuffered TSB without glucose or buffered pH 5.5 or TSB pH 5.5 supplemented with palmitic acid (C16:0), palmitoleic acid (C16:1), or both fatty acids combined (C16:0 + C16:1) as indicated. Biofilm biomass was measured at 48-hours using crystal violet staining. Data represent mean ± SEM performed in triplicate. Statistical significance was measured using one-way ANOVA with Tukey’s multiple comparisons test, ****p<0.0001, ns; not significant.

### Skin-derived chemical cues elicit differentially affect extracellular protease profiles in *S. aureus* and *S. epidermidis*

*S. aureus* and *S. epidermidis* have orthologous serine proteases SspA (V8 protease) and Esp respectively. Previous studies indicated that *S. epidermidis* Esp can eradicate *S. aureus* nasal colonization through disruption of biofilm formation, while SspA of *S. aureus* has been implicated in promoting itchy skin in atopic dermatitis (29, 30). Moreover, we found that expression of SspA in *S. aureus* is induced in response to acidic pH and antimicrobial unsaturated free fatty acids that would be encountered on human skin (20). We therefore conducted experiments to determine if this was also true for *S. epidermidis*. Since our previous work was conducted using TSB with glucose, we first assessed secreted protein and protease production in unbuffered TSB with glucose. Under this condition, *S. epidermidis* exhibited abundant production of a ∼26 kDa peptide that was identified by mass spectrometry as Esp (Fig S6), accompanied by detection of protease activity in a casein zymogram (Fig 5A), whereas no protease production was evident in *S. aureus* USA300 under the same condition. In TSB with no glucose, the 26 kDa peptide corresponding to Esp in *S. epidermidis* was no longer evident, and no protease activity was detected in the zymogram assay for either *S. aureus* or *S. epidermidis*.

**Fig 5:**
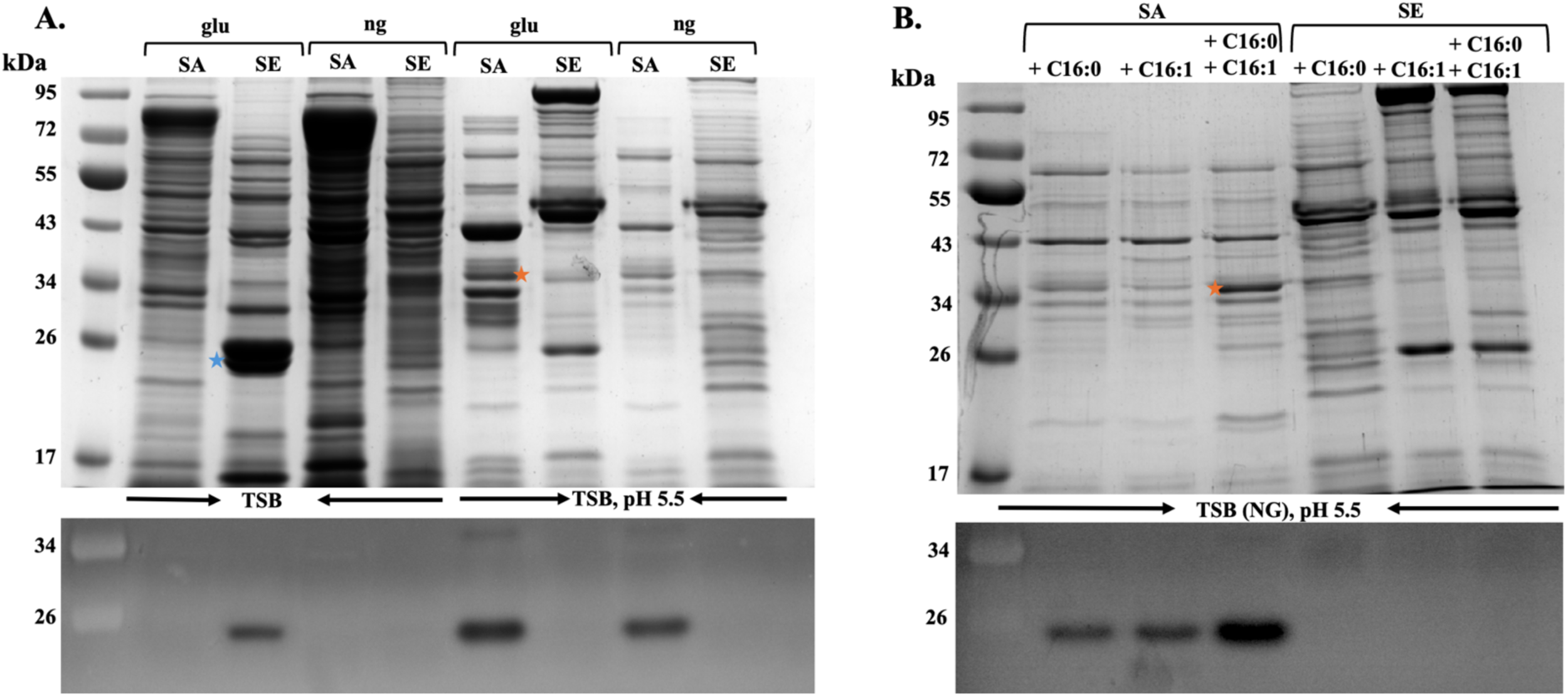
Influence of glucose and acidic pH (A), or acidic pH and C16 fatty acids (B) on secreted protein profiles (top panels) and protease production (bottom panels) by *S. aureus* USA300 (SA) and *S. epidermidis* M23864:W2 (SE). (A), Cultures of *S. aureus* (SA) or *S. epidermidis* (SE) were grown in unbuffered TSB or TSB pH 5.5 as indicated. Presence or absence of glucose is indicated by (glu) or (ng) respectively above each lane. (B), cultures were grown in glucose free TSB buffered at pH 5.5 supplemented with C16:0 (500 µM), C16:1 (25 µM) or both C16:0 and C16:1 as indicated. After growth for 20 hours and determination of OD_600_, cultures were centrifuged and samples of cell free culture supernatants were processed for analysis of secreted proteins by SDS-PAGE and Coomassie staining of secreted proteins (top panels), or zymography for detection of protease (bottom panels). For secreted protein profiles, each lane contains TCA-precipitated protein from 2.5 OD_600_ units of culture supernatant, while for zymograms a volume of culture supernatant equivalent to 0.075 OD_600_ units was applied to each lane. Star indicates the position of signature proteins, mature SspA (orange stars) and Esp serine protease (blue stars).

Although *S. aureus* USA300 did not produce detectable protease when cultured in TSB with glucose, when this medium was buffered at pH 5.5 SspA production was stimulated (Fig 5A) as indicated by a cluster of ∼ 36 kDa polypeptides (Fig S6), which is characteristic of intermediates in processing of the N-terminal propeptide of proSspA (41), and the more abundant lower mass protein was confirmed as SspA through mass spectrometry (Fig. S6). Production of SspA by *S. aureus* was maintained in TSB pH 5.5 that lacked supplemental glucose, whereas no proteolytic activity was evident in *S. epidermidis* in TSB pH 5.5 irrespective of the inclusion or absence of glucose. At pH 5.5 with no glucose, *S. aureus* maintained SspA production when supplemented with either 500 µM palmitic acid or 25 µM palmitoleic acid, and the presence of both fatty acids stimulated SspA production, whereas *S. epidermidis* again failed to produce Esp when supplemented with these fatty acids alone or in combination (Fig 5B). We conducted additional tests to assess protease production in a cross section of genetically diverse *S. epidermidis* isolates recovered from human skin (42) and overall, the data are consistent with a requirement for glucose to produce Esp in *S. epidermidis*, whereas acidic pH represses protease production irrespective of the presence or absence of glucose, and exposure to C16 fatty acids was not sufficient to induce protease production (Fig S3). These findings underscore a distinct regulatory divergence between *S. aureus* and *S. epidermidis* in protease production where *S. aureus* actively responds to skin chemical cues through protease upregulation, while Esp production in *S. epidermidis* is restricted under the same conditions.

### *S. epidermidis* GraRS two-component system functionally complements *S. aureus graRS* mutants but exhibits reduced efficiency under fatty acid stress

The GraRS two-component regulatory system in *S. aureus* is essential for resistance to skin-associated stressors, including acidic pH, antimicrobial peptides, and fatty acids (20, 21, 43). Intriguingly, the GraS sensor kinase activity was needed for production of SspA protease at acidic pH in *S. aureus* (20). We therefore queried whether differential growth and protease expression in response to environmental cues could be accounted for through differences in signaling through GraS, since there is 70% sequence identify for GraS between *S. aureus* and *S. epidermidis*, compared to 91% for GraR. In TSB pH 5.5, there were no discernible differences in growth between USA300 and its isogenic *graS* and *graR* deletion mutants (Fig S4A, C). However, as we reported previously (20), USA300Δ*graS* exhibited significantly enhanced binding of cytochrome C relative to USA300 (Fig 6A), reflecting loss of signaling through GraS and failure to express MprF which reduces the negative cell surface charge through lysine modification of phosphatidylglycerol. Strikingly, cell surface charge was equally restored through complementation with either *graS* (SA) or *graS* (SE) indicating that the orthologous sensor kinases are functionally interchangeable in response to acidic pH alone. However, a different story emerged when growth at pH 5.5 in presence of 15 µM palmitoleic acid was employed as the basis of comparison. Under this condition, growth of USA300Δ*graS* was strongly impaired, and normal growth was restored by complementing with *graS* (SA), whereas *graS* (SE) was significantly less effective (Fig 6B). Nevertheless, when growth was extended to 24h, *graS* (SE) showed some restoration of growth (Fig S4B).

**Fig 6:**
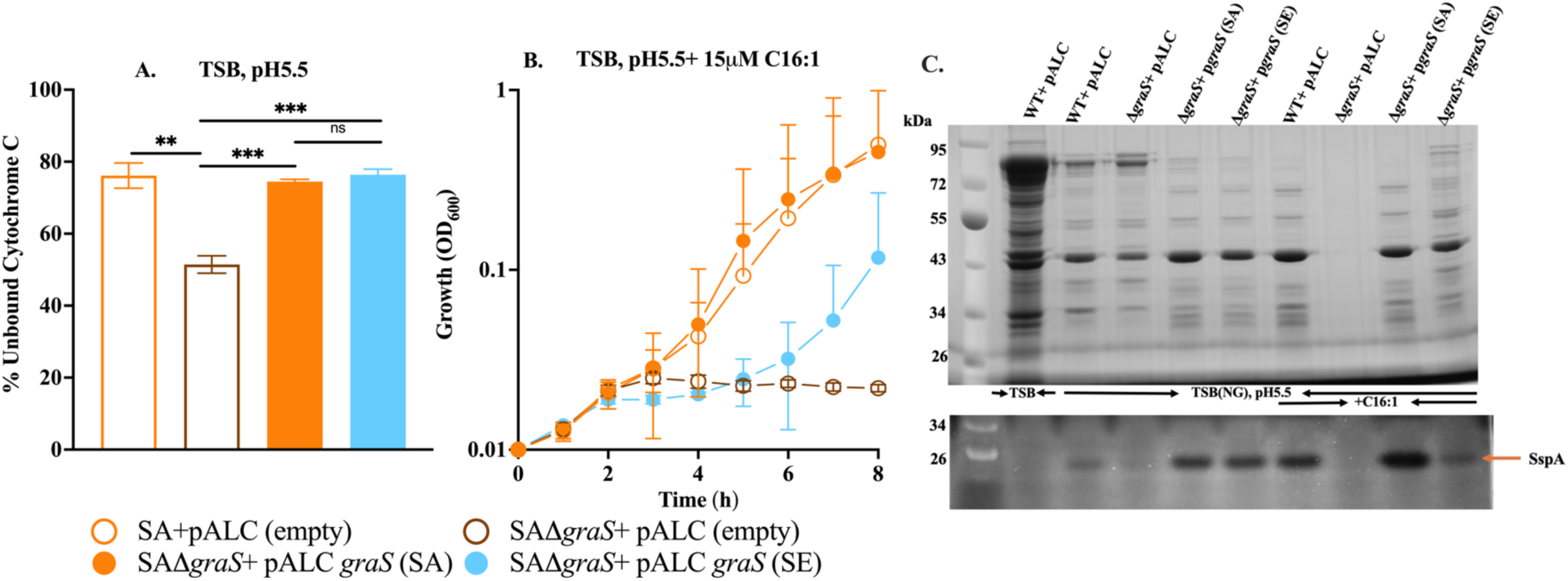
Complementation of *S. epidermidis graS* restores cell surface charge (A), growth (B) and protease production (C) in *S. aureus* D*graS* mutant. (A) Cell surface charge was assessed using the cytochrome C binding assay in the TSB media without glucose buffered pH5.5 (TSB (NG), pH5.5). (B) Growth kinetics of the above strains cultured in TSB (NG), pH5.5 and supplemented with 15 µM palmitoleic acid (C16:1). Growth (OD_600_) was monitored over 8 hours. (C) Secreted protein and protease profiles of strains grown for 24 h in TSB (with glucose, unbuffered) or without glucose pH 5.5 with or without 15 µM C16:1. Culture supernatants were processed for analysis of secreted proteins by SDS-PAGE and Coomassie staining (top panels), or zymography for detection of protease (bottom panels). For SDS-PAGE, each lane contains TCA-precipitated proteins from 2.5 OD_600_ units of culture; for zymography, supernatant volume equivalent to 0.075 OD_600_ units was loaded. The arrow (orange) indicates the position of the mature serine protease SspA. Data represent mean ± SEM from three biological replicates. Statistical significance was assessed using one-way ANOVA with Tukey’s multiple-comparison test (**p < 0.01, ***p < 0.001, ns = not significant).

Similar outcomes were noted when protease production was assessed (Fig 6C). Growth at pH 5.5 alone stimulated production of SspA in wild type USA300 compared to unbuffered TSB, and diminished protease production was evident in USA300Δ*graS*. Moreover, just as *graS* (SA) and *graS* (SE) were equally effective in restoring cell surface charge during growth at pH 5.5, complementation with either sensor kinase promoted enhanced production of SspA. Supplementation of TSB pH 5.5 with 15 µM palmitoleic acid strongly enhanced production of SspA in USA300 compared to pH 5.5 alone. With these combined stressors, production of SspA and secreted proteins in USA300Δ*graS* could not be assessed, since growth was strongly impaired. However, when USA300Δ*graS* was complemented with *graS* (SA), protease production exceeded that of wild type USA300, while *graS* (SE) was once again less effective as also noted in growth analyses. Similar results were obtained when functional interchangeability was assessed by complementing Δ*graR* with either *graR* (SA) or *graR* (SE) (Fig S5). Both homologs successfully restored growth, protease activity, and surface charge in the Δ*graR* mutant, although as with GraS, the *graR* (SE) complementation was less efficient under combined acidic pH and fatty acid stress. Collectively, these data demonstrate that the *S. epidermidis* GraRS system can functionally substitute for its *S. aureus* counterpart in regulating growth, protease production, and membrane surface charge under acidic conditions. However, its reduced efficiency under fatty acid stress suggests species-specific optimization of this regulatory pathway. Consequently, it is plausible that species-specific promoter architecture or regulator-binding motifs for SspA and Esp underlie differential target gene regulation by GraRS across these species.

### Acidic pH and fatty acid induce *sspA* but not *esp* promoter activity in *S. aureus*

In view of our finding that *graS* (SE) and *graR* (SE) could only partially restore growth and protease production in *S. aureus ΔgraS and ΔgraR* mutants under combined exposure to acidic pH and C16 fatty acids, we speculated that the differential protease expression of orthologous proteases in these species could be attributed to the promoter elements and interaction with regulatory proteins. As a first step towards addressing this, we constructed *sspA::lux* and *esp::lux* reporters where luciferase expression is driven from the respective promoter elements of these proteases. In TSB with no glucose at pH 5.5, *sspA::lux* exhibited peak expression at ∼ 5h (Fig 7A) and expression was enhanced with 500 µM palmitic acid (Fig 7B), while *esp::lux* was inactive under identical conditions. In unbuffered TSB with glucose, under which condition we did not detect SspA activity on zymograms, *sspA::lux* was considerably less active compared to TSB pH 5.5, but surprisingly this condition was more permissive to *esp::lux* activity, which was comparable to *sspA::lux* from 5-8h of growth (Fig 7C). Therefore, the *sspA::lux* and *esp::lux* reporters follow the same expression profiles in *S. aureus* as was evident from detection of SspA and Esp production under different growth conditions for either microbe.

**Fig 7:**
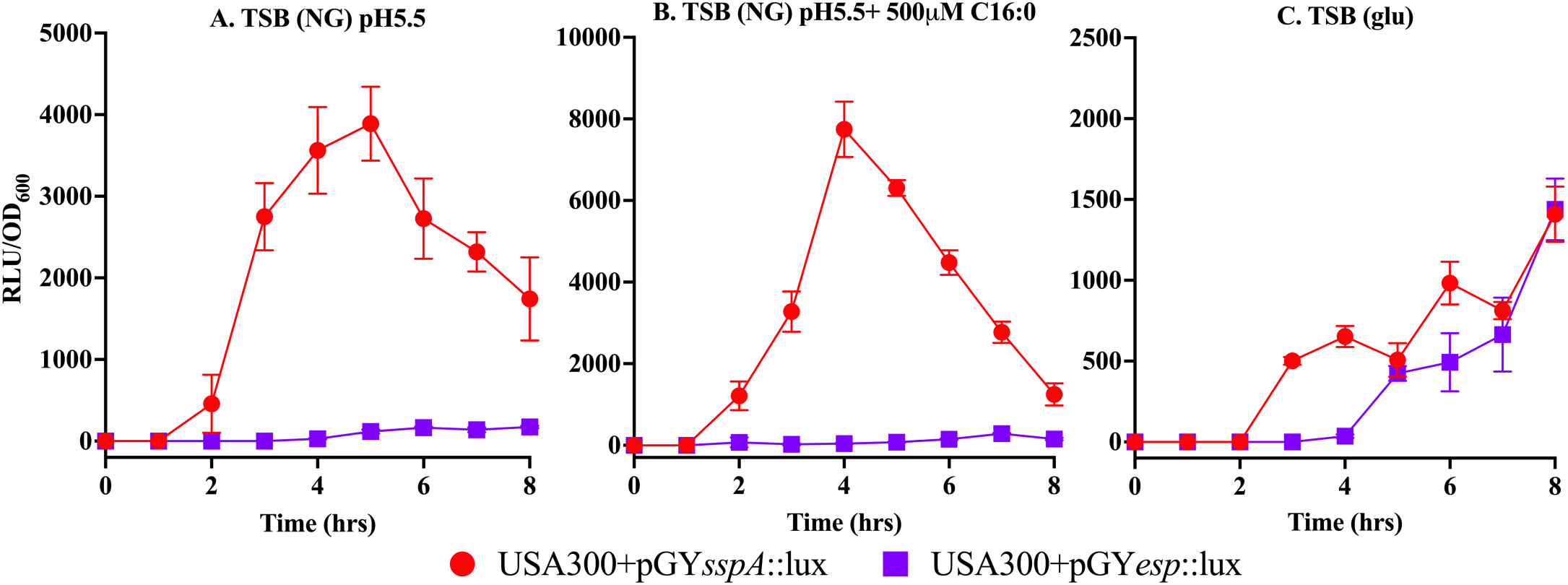
Differential promoter activity of *sspA* and *esp* in *S. aureus* USA300 under skin like conditions. *S. aureus* USA300 strains carrying the lux reporter plasmid with either the *sspA* promoter (pGY*sspA*::lux) or the *esp* promoter (pGY*esp*::lux) were grown in different conditions: (A) TSB without glucose buffered pH5.5 [TSB(NG), pH5.5] alone or (B) with 500mM palmitic acid (C16:0) and (C) TSB with glucose [TSB (glu)] at unbuffered condition. Bioluminescence was measured at indicated time points in the presence of decanal and normalized to cell density (RLU/OD_600_). The *sspA* promoter exhibited strong induction under acidic conditions and in the presence of palmitic acid (C16:0), with maximal activity at 4–6 hours, while the *esp* promoter showed minimal or no induction under the same conditions. Data represent mean ± SEM from triplicate flasks

## DISCUSSION

Adaptation to skin’s chemical stressors is a key determinant of microbial survival and competitiveness on the skin surface (44), and our study provides important new insight into how *S. aureus* and *S. epidermidis*, two closely related Staphylococci, respond to skin-derived chemical signals. At pH 5.5 with unsaturated palmitoleic acid, *S. aureus* growth was significantly impaired whereas *S. epidermidis* was largely unaffected, indicating a higher intrinsic tolerance of the commensal to low pH and C16 unsaturated fatty acids encountered on human skin. This aligns with previous findings that *S. epidermidis* has a higher MIC for unsaturated sapienic acid compared to *S. aureus* (45), as also noted here with its isomer palmitoleic acid. Conversely, *S. aureus* exhibited a growth advantage over *S. epidermidis* in presence of palmitic acid alone, consistent with it having a functional *fadXDEBA* locus for metabolism of palmitic acid through β-oxidation, whereas this pathway is incomplete in *S. epidermidis* (5). Furthermore, although palmitoleic acid was more toxic to *S. aureus* at acidic pH compared to *S. epidermidis*, this toxicity was mitigated in the presence of palmitic acid. Collectively, these findings underscore the role of host-derived fatty acids in modulating Staphylococcal colonization, while shedding new insight into how *S. aureus* and *S. epidermidis* differentially respond to the combined acidic pH and C16 fatty acids that are abundant on human skin, as manifested through our analysis of phospholipid composition, biofilm formation and protease production.

The mature Esp glutamyl endopeptidase of *S. epidermidis* shares 51% identity and 70% similarity with the orthologous SspA of *S. aureus*. Consistent with a recent report in which SspA was found to promote itchy skin in atopic dermatitis through proteolytic processing of the PAR1 receptor (29), we find that SspA production is stimulated in *S. aureus* by acidic pH and unsaturated fatty acids that would be encountered on human skin. However, these same conditions restricted production of Esp by *S. epidermidis*, and we confirmed this with additional human commensal strains. Others noted that Esp activates pro-IL1β which would also be proinflammatory (46), and inconsistent with *S. epidermidis* existence as a commensal on human skin. It is not known whether *S. aureus* SspA also exhibits this activity. However, it is noteworthy that *Streptococcus pyogenes* colonization of human skin is restricted by the SpeB cysteine protease activation of Gasdermin A, which initiates pyroptosis in keratinocytes (47), and we have recently described a similar activity for the Staphopain A (ScpA) protease of *S. aureus* (48). As such, our observations of differential expression of the orthologous Esp and SspA glutamyl endopeptidases of *S. epidermidis* and *S. aureus* in response to acidic pH and C16 fatty acids encountered on human skin are consistent with their respective commensal and pathogenic phenotypes.

Previous work implicated *S. epidermidis* Esp in controlling *S. aureus* colonization of humans through disruption of biofilm formation (30). However, this role is inconsistent with our finding that acidic pH and C16 fatty acids restrict production of Esp, while stimulating expression of SspA. Accordingly, we did observe that biofilm production was differentially influenced by these conditions. Notably, biofilm formation by both bacteria was stimulated with combined exposure to acidic pH and 500 µM palmitic acid. However, for *S. aureus* where a combination of palmitic and palmitoleic acid stimulates SspA production, the stimulatory activity of palmitic acid was abrogated by palmitoleic acid, while enhanced biofilm formation was maintained in *S. epidermidis* under this same condition. These findings are consistent with *S. epidermidis* biofilm formation being optimized to signals encountered on human skin, and insofar as protease production is counterproductive to biofilm formation, our data on differential protease expression and biofilm formation are also consistent with the pathogen and commensal phenotypes of *S. aureus* and *S. epidermidis* respectively.

Although there is considerable variability in the composition of biofilm matrices, including nucleic acid, protein and polysaccharide components, we can exclude the intercellular polysaccharide adhesin Ica as a mechanism of biofilm formation for *S. epidermidis* M23864:W2, since like many other commensal *S. epidermidis* strains, it lacks the *ica* locus which is prevalent among *S. epidermidis* isolates from nosocomial infections (49, 50), and the *ica* locus is also highly prevalent among *S. aureus* clinical isolates (51). It is therefore likely that biofilm formation in *S. epidermidis* commensal strains is reliant on documented cell surface proteins (52), most of which have orthologs in *S. aureus*. Therefore, evolution of commensal *S. epidermidis* to exhibit low protease production in response to combined acidic pH and host derived fatty acids encountered on skin is consistent with a requirement for proteinaceous biofilms in maintenance of colonization. However, it is less clear how *S. aureus* and *S. epidermidis* achieve differential expression of conserved glutamyl endopeptidases in response to common environmental signals, given the maintenance of orthologous regulators in both microbes, including activation through quorum sensing by the accessory gene regulator *agr* locus, and repression through the *sarA* regulator (53). In view of these considerations, we focused our attention on the GraRS two component systems.

GraRS was discovered independently in *S. aureus* and *S. epidermis* in relation to its role in resistance to cationic antimicrobial peptides, which in both cases involves signaling through GraS to phosphorylate GraR and promote expression of a phosphatidylglycerol lysyl-transferase MprF which adds a positive charge to the phospholipid, thereby repelling cationic antimicrobial peptides (17, 20). However, our work expanded the role of GraS in *S. aureus* through our discoveries that signaling is also activated by acidic pH, under which condition GraS promotes expression of SspA and enhanced resistance to unsaturated fatty acids. It was therefore surprising that during growth at acidic pH alone GraS (SE) and GraR (SE) were equally effective as *S. aureus* GraSR in restoring cell surface positive charge and SspA expression in *S. aureus* USA300*ΔgraS* and USA300*ΔgraR* mutants. Although this correlation breaks down during combined exposure to acidic pH and palmitoleic acid, our data reveal that at acidic pH alone, the functions of GraRS in *S. aureus* and *S. epidermidis* are interchangeable with respect to maintenance of cell surface charge, even though *S. epidermidis* did not produce Esp at acidic pH. A plausible explanation is that GraS is an intramembrane sensor kinase that responds to membrane perturbation (54, 55). As such, our data have revealed for the first time significant differences in membrane phospholipid composition when *S. aureus* and *S. epidermidis* are cultured at acidic pH alone, or acidic pH combined with C16 fatty acids that would be encountered on human skin.

GraS is an intramembrane sensor kinase that signals in response to membrane perturbation, and as with the SaeS sensor kinase that belongs to this same family and modulates virulence factor expression in *S. aureus* (56), the extracellular sensor domain of GraS is comprised of just nine amino acids flanked by transmembrane spanning helices (57). As such, the minimal sensor domain has been described as a molecular tripwire that is activated by membrane perturbation (56, 58). We previously speculated that reduced membrane curvature imposed by a reduction in the repulsive forces between surface exposed polar phospholipid head groups could be one factor in activation of signaling through GraS at acidic pH, and changes in phospholipid composition represents another factor. A striking difference during growth at acidic pH was that *S. epidermidis* PG consisted of 79% PG35 (C15 + C20) and PG33 (C15 + C18) compared to 55% in *S. aureus*, and this difference in PG species abundance in *S. aureus* was compensated by a higher content of PG31:0 (C15 + C16) fatty acids, such that at acidic pH alone, *S. epidermidis* PG had a higher content of longer C18 and C20 straight chain fatty acids. This follows a pattern where in other bacteria at low pH extremity, the acyl chains of phospholipid are lengthened to restrict passage of protons across a thicker membrane (59). Therefore, while the signaling mechanism through GraS in each microbe may be functionally conserved, the bacteria exhibit significant differences in phospholipid composition at acidic pH, which may influence the ability of GraS (SE) to stimulate SspA production in *S. aureus* during growth at pH 5.5, even though this does not occur in *S. epidermidis*. Importantly, the similarly functioning SaeS regulator binds to both cardiolipin and phosphatidylglycerol (60), consistent with our hypothesis that differences in phospholipid composition may contribute to differential signaling through GraS in *S. aureus* and *S. epidermidis* under identical growth conditions.

Consistent with being better adapted to growth on human skin, *S. epidermidis* exhibited a greater capacity to directly incorporate the C16 fatty acids that are most abundant on human skin into its phospholipid. Remarkably, when cultured at pH 5.5 with 500 µM palmitic acid, *S. epidermidis* phospholipid contained 32% PG32:0 with palmitic acid at both the sn1 and sn2 positions, compared to less than 4% in *S. aureus*. Under this same condition, *S. aureus* had significantly elevated PG35:0 (C15 + C20) compared to *S. epidermidis*, consistent with previous findings that it has a stringent preference for incorporation of branched chain C15 fatty acid at the sn2 position of PG, while exogenous palmitic acid is extended to C20 via FASII prior to incorporation into PG at sn1 through the combined action of FakA, PlsX and the PlsC acyltransferase (23, 33). It is noteworthy that in *E. coli*, PlsC accepts both acyl-phosphate and acyl-ACP as substrates (61), whereas current literature indicates that in Gram-positive bacteria, PlsC is restricted to an acyl-ACP substrate. However, this conclusion is based on research conducted with *Streptococcus pneumoniae* and *S. aureus* (33). Of interest, PlsC of *S. aureus* and *S. epidermidis* share only 81% identity, compared to 89% for PlsY. Whether this greater sequence divergence of PlsC is sufficient to account for the differential capacity of *S. epidermidis* to incorporate C16 fatty acids into phospholipid remains to be determined. However, as noted for PlsC in *E. coli*, it is feasible that the ability to utilize both C16 acyl-phosphate and C16 acyl-ACP as substrate is a phenotype that is conducive to commensalism.

A last consideration for the differential responses of *S. aureus* and *S. epidermidis* towards common environmental signals encountered on human skin is variation in the promoter elements of genes with orthologous functions, of which we have focused on the orthologous Esp and SspA proteases. Major known regulators of the proteolytic phenotype in *S. aureus* which have orthologs in *S. epidermidis* include the alternative Sigma Factor SigB, the SarA repressor, the accessory gene regulator locus Agr, and GraSR (62–64). Strikingly, even though GraS (SE) was able to functionally complement GraS (SA) with respect to production of SspA at acidic pH, the *S. epidermidis esp::lux* reporter did not respond to acidic pH when expressed in *S. aureus*, indicating that its promoter element lacked the necessary structural details to interact with the relevant regulators in *S. aureus* needed for expression at acidic pH. Moreover, when *esp::lux* activity was assessed in *S. aureus*, its expression was optimal under conditions where optimal Esp production was also evident in *S. epidermidis*, such that expression was highest in TSB with glucose and not evident in TSB pH 5.5 or pH 5.5 with palmitic acid, under which condition *sspA::lux* was highest. Therefore although *S. aureus* and *S. epidermidis* share several orthologous regulators that influence protease expression, the promoter elements of Esp and SspA appear to be sufficiently divergent to permit their differential expression on exposure to acidic pH and C16 fatty acids that are encountered on human skin, and it is further feasible that as yet unidentified regulators that may be unique to either microbe may contribute to this differential expression.

In summary, our study was initiated in response to reports that the Esp protease of *S. epidermidis* could modulate *S. aureus* colonization of human skin and anterior nares through disruption of biofilm formation (30). Although we recognize that additional environmental and nutritional cues unique to the ecological niche on human skin may be permissive to Esp production *in vivo*, the body of work presented here is supportive of Esp expression in *S. epidermidis* being restricted as a means of maintaining a proteinaceous biofilm matrix and avoiding deleterious pro-inflammatory consequences of protease expression, while under the same conditions, expression of SspA by pathogenic *S. aureus* is favoured. These differential phenotypes may in part be attributed to differences in phospholipid composition and metabolism of C16 fatty acids whereby *S. aureus* has a functional beta oxidation pathway and restricts incorporation of C16 fatty acids into phospholipid, whereas this pathway is incomplete in *S. epidermidis* which has the capacity to directly incorporate palmitic acid into phospholipid. Cumulatively, these findings represent significant new insight into how pathogenic and commensal phenotypes of *S. aureus* and *S. epidermidis* are modulated by the common signals to which they are exposed on human skin.

## MATERIALS AND METHODS

### Bacterial strain and growth conditions

Bacteria and plasmids that were used or constructed in this study are listed in Table S2 in the supplemental material. *S. aureus* cultures were maintained as frozen stocks (-80°C) in 20% glycerol and streaked on TSB agar (TSA) when required. Tryptic soy broth (TSB) containing 2.5 g L^−1^ glucose (∼14 mM) or TSB without glucose were supplied by Bacto. TSB or TSA were supplemented when needed, with 10 μg mL^−1^ erythromycin or chloramphenicol for propagation of strains bearing resistance markers. Where indicated, TSB or TSA was supplemented by addition of 0.1 M morpholineethanesulfonic acid (MES) buffer (Bio Can Scientific) pH 5.5 prior to autoclaving. To supplement media with palmitic acid, a 100 mM stock concentration of palmitic acid was first prepared in 70% ethanol and then diluted into TSB 0.1% DMSO to achieve the desired concentration. To supplement with palmitoleic acid, a 5mM stock concentration was first prepared in TSB containing 0.1% dimethyl sulfoxide (DMSO) and then diluted into TSB plus 0.1% DMSO to achieve the desired concentration. Palmitic acid (hexadecanoic acid; C16:0) was purchased from Sigma, and palmitoleic acid (cis-9-hexadecenoic acid; 16:1) was purchased from Cayman Chemicals. *E. coli* strains were grown on LB agar or LB broth supplemented with 100 μg mL^−1^ ampicillin. Unless otherwise stated, all cultures were grown at 37°C, and liquid cultures were incubated on an orbital shaking platform at 220 rpm. Precultures were grown in 3 mL of media in a 13-mL polypropylene tube, whereas experimental cultures were grown in 25 mL of media within 125-mL-capacity flasks, to allow for proper aeration while shaking.

### Strains and plasmid construction

Genetic manipulation of *S. aureus* was conducted according to established guidelines and as described in previous work (20, 65, 66). Restriction enzymes and T4 DNA ligase were purchased from New England Biolabs, Taq polymerase from GenScript, PCR cleanup and plasmid preparation kits were from Geneaid and nucleotide primers (Table S3) from Integrated DNA Technologies. All recombinant plasmids were initially constructed in *E. coli* DH5α. The integrity of plasmids was confirmed through nucleotide sequencing (Plasmidsaurus) prior to electroporation into USA300 or isogenic derivatives, using *S. aureus* RN4220 as an intermediate host. Primer sequences used for PCR amplification of gene segments for plasmid construction are defined in Table S3 in the supplemental material.

Deletion of *graR* was performed using the temperature sensitive pKOR mutagenesis plasmid and procedure as previously described (67). Briefly, ∼1 kb upstream and downstream regions of *graR* were amplified through PCR. The primers *graR*-UP-attB1 with *graR*-UP-SacII were used to construct the upstream amplicon, and the primers *graR*-DW-attB2 with *graR*-DW-SacII were used to construct downstream amplicon, as outlined in Table S3. Upstream and downstream amplicons were digested with *SacII* and ligated together using T4 DNA ligase. Ligation products were then recombined into the parental pKOR plasmid using Gateway® BP Clonase® (Thermo Fisher Scientific). The resulting pKOR::Δ*graR,* deletion plasmid was passage through *E. coli* DH5α and *S. aureus* RN4220, before electroporation into *S. aureus* USA300. Through homologous recombination, the *graR* gene was excised and replaced with an in-frame deletion as previously described (67). Successful deletion of *graR* was confirmed through PCR and DNA sequencing using the primers *graR-*DEL-UP and *graR*-DEL-DW, detailed in Table S3. Plasmid pALC2073, which provides a basal level of gene expression from the Pxyl/tetO promoter and a stronger inducible level of expression with anhydrotetracycline (68, 69), was used for ectopic expression of *graS* and *graR* from *S. aureus* (SA) or *S. epidermidis* (SE). Plasmid pGYlux was used to quantify *sspA* and *esp* promoter (70). Primer pairs *sspA-p*-F/*sspA-p*-R and *esp-p*-F/*esp-p-R* were used to amplify the prompter region of *sspA and esp* respectively, from *S. aureus* USA300 LAC genomic DNA and *S. epidermidis* M23864:W2 genomic DNA. The resulting amplicons, and the pGYlux plasmid, were digested with *BamHI*-HF and *SalI*-HF, and ligated together with T4 DNA ligase, to construct pGY*sspA*::lux and pGY*esp*::lux respectively.

### Growth and MIC assay

For growth analyses, cultures of *S. aureus* or *S. epidermidis* were prepared by inoculating 3ml of TSB in a 13-ml polypropylene tube containing antibiotic as required and grown overnight for 16 h. After determining the optical density at 600 nm (OD_600_), aliquots were subcultured into 125-ml-capacity flasks containing 25 ml of TSB or TSB modified by addition of buffer and fatty acid to achieve an initial OD_600_ of 0.01. Growth (OD_600_) was monitored at hourly intervals. For MIC assays, inoculum cultures were grown to mid-exponential phase in flasks as for growth assays and then subcultured at an OD_600_ of 0.01 into triplicate 20-by 150-mm glass culture tubes containing 3 ml of medium supplemented with 0.1% DMSO and designated concentrations of palmitoleic acid. Cultures were incubated at 37°C with vigorous shaking, and OD_600_ values were determined after 24 h.

### Phospholipid analysis

For lipid extraction, a modified Bligh and Dyer method was used (5, 71). Briefly, triplicate cultures of *S. aureus* and *S. epidermidis* were grown in TSB without glucose buffered pH5.5 alone or supplemented with palmitic acid or with palmitic and palmitoleic acid for 4 to 6 h. Then approximately 2′10^8^CFUs of culture was pelleted and washed with 10 mL of PBS three times. The washed samples were then resuspended in 70% methanol (w/v) and transferred to a glass tube with Teflon lined cap. Each sample was supplemented with 100ng of PG-d7 (15:0/18:1) internal phosphatidylglycerol standard, followed by addition of 5 mL methyl tert-butyl ether. After mixing, the tubes were capped and incubated for 30 minutes at room temperature in a water bath sonicator. After adding 1.25 mL of water, mixing and centrifugation, the clear upper layer was collected into a fresh glass tube. The extraction process was repeated twice more, and the combined organic extract was evaporated under nitrogen. Each dried sample was dissolved in 40 µl of solvent A (Water: Acetonitrile:ammonium acetate 90:10:0.1) and 60 µl of solvent B (5:95:0.1).

HPLC was performed on a Prominence XR system (Shimadzu) using a Targa C8 (2x20 mm, 5µ, Higgins Analytical) column. The mobile phase consisted of a gradient between buffer A (water-acetonitrile 90:10 v/v) and buffer B (water-acetonitrile 5:95 v/v), both containing 0.1% ammonium acetate pH 8.0. Lipids were separated over a gradient of 0 to 60% buffer B over 1 minute, and then 60 to 100% buffer B over 9 minutes, at a flow rate of 0.5 mL/min. The HPLC eluate was fed directly to the ESI source of a QTRAP5500 mass analyzer (SCIEX) in the negative ion mode with the following conditions: Curtain gas, GS1, and GS2: 35 psi, Temperature: 600 °C, Ion Spray Voltage: -2500 V, Collision gas: low, Declustering Potential: -60 V, and Entrance Potential: -7 V. The eluate was monitored by the Multiple Reaction Monitoring (MRM) method to detect unique molecular ion–daughter ion combinations. All possible PGs are from m/z 600-1000 Da with product ion m/z 153 (negative ion mode). The MRM is scheduled to monitor each transition for 120 s around the established retention time for each lipid mediator. Optimized Collisional Energy (-40 eV) and Collision Cell Exit Potentials (– 10 V) are used for each MRM transition. Mass spectra for each detected lipid mediator were recorded to MRM transition and retention time match with the standard. Data were collected using Analyst 1.6.2 software, and the MRM transition chromatograms quantified by SCIEX software. The internal standard signals in each chromatogram were used for normalization for recovery as well as relative quantitation of each analyte.

### Laurdan membrane fluidity assay

Laurdan membrane fluidity assay was performed as previously described (72). Briefly, overnight cultures of *S. aureus* or *S. epidermidis* were inoculated into 125 mL flasks containing 25 mL of TSB without glucose pH 5.5 alone or with supplemental fatty acids as indicated, to achieve an initial OD_600_ of 0.01, and grown to mid log-phase at 37°C with shaking. After normalizing to OD_600_ of 0.4 with PBS, 1 mL of culture was centrifuged for 8 min at 10,000 xg and the cell pellet was washed twice with 1 mL of PBS, after which the cell pellet was resuspended in 1 mL of 10 µM Laurdan in Laurdan buffer (Table S4). After incubation in the dark for 10 min at 37°C, the cell suspension was again centrifuged, washed in 1 mL of PBS, and finally resuspended in 200 mL of fresh Laurdan buffer. The cell suspensions were then transferred to 96-well black microtiter plates (Agilent) for measurement of fluorescence with excitation at 350 nm and emission at 435 nm and 490 nm, using a Cytation 5 multimode plate reader (Agilent Instruments). The Laurdan GP-ratio was calculated from the emission outputs at different wavelengths using the formula (435-490)/(435+490) as described previously (73, 74).

### Biofilm measurement

Biofilm accumulation during growth in microtiter plates was quantified using crystal violet stain as previously described (75, 76). Briefly, overnight cultures of *S. aureus* and *S. epidermidis* grown in TSB with glucose were diluted to OD_600_ of 0.01 in different media including TSB with no glucose, TSB with no glucose buffered at pH 5.5, and TSB no glucose pH 5.5 supplemented with either 500 µM palmitic acid, 25 µM palmitoleic acid, or with both 500 µM palmitic and 25 µM palmitoleic acid. Triplicate aliquots of 200 µl were then transferred into 96 well polystyrene microtiter plates (Fisher scientific, Costar^TM^ 96-well flat bottom) which were incubated statically for 24 h or 48 h at 37°C. After incubation, planktonic cells were aspirated, and the plates were gently twice submerged in water followed by shaking to remove unattached cells. The remaining biofilm was then fixed by incubation with 150 mL of methanol for 15 min. After washing the fixed biofilm with water, the plates were allowed to air dry at room temperature followed by addition of 150 mL of 0.5% crystal violet stain and incubation at room temperature for 20 min. After gentle washing 4 times with water as described above the stained biofilms were allowed to air dry, followed by addition of 150 mL of 30% glacial acetic acid to each well. After incubation for 15 min at room temperature to solubilize the crystal violet stain, the liquid was transferred to a fresh flat bottom microtiter plate to measure A_550_ using 30% glacial acetic acid as the blank.

### SDS-PAGE and Zymography

For SDS-PAGE analysis of secreted protein profiles, *S. aureus* cultures were grown for 20 h, and proteins in cell-free culture supernatant were precipitated by mixing with equal volumes of ice-cold 20% trichloroacetic acid (TCA), washed in ice-cold 70% ethanol, and then air dried prior to dissolving in SDS-PAGE reducing buffer as described previously (41). Protein equivalent to 2.5 OD_600_ units of culture supernatant was then loaded for protein separation on a 10% acrylamide gel using the Laemmli buffer system (77) and after electrophoresis, proteins were stained using Coomassie blue. For detection of protease activity by zymogram assay, the resolving gel was copolymerized with 1 mg/ml casein and protein equivalent to 0.075 OD_600_ unit was applied to each lane. Details on sample processing, electrophoresis, and zymogram development are as described previously (78).

### Trypsin digestion and mass spectrometry

Where indicated, Coomassie stained proteins bands from SDS-PAGE were excised with a sharp scalpel and submitted to the University of Toronto Sick Kids Hospital SPARC Biocentre for trypsin digestion and protein identification by mass spectrometry. Tryptic peptides were eluted from the gel slices in acetonitrile, and after drying under vacuum, the samples were desalted on a C18 ziptip, and resuspended in 0.1% formic acid. Mass spectrometry (LC-MS/MS) was performed using the EASY-nLC 1200 nano-LC system and a Thermo Scientific Q Exactive HF-X mass spectrometer (Fisher Scientific Company, Ottawa, ON, Canada). Protein identification and quantification were performed using Proteome Discoverer (MS Amanda and Sequest HT) and Scaffold (X! Tandem), which were setup to search the NCBI nr database of *S. aureus* protein sequences.

### Cytochrome C binding assay

Cell surface charge was measured as a function of cytochrome c binding as previously described (20, 79). Briefly, bacterial cultures were grown to an OD_600_ of 0.5 before being washed twice in morpholinepropanesulfonic acid (MOPS) buffer (20 mM, pH 7.0). Cells were resuspended to an OD_600_ of 7.0 in MOPS buffer, and 360 mL aliquots were mixed with 40 mL of bovine cytochrome c (Sigma) to a final concentration of 0.5 mg/ml. Samples were incubated for 15 min at 37°C, followed by centrifugation at 6,000 g for 8 min, and the remaining unbound cytochrome c was quantified by measuring absorbance at 530 nm (A_530_) relative to a MOPS buffer blank containing 0.5 mg/ml cytochrome C.

### Bioluminescence reporter assay

Overnight cultures of USA300 pGY*sspA*::lux or USA300 pGY*esp*::lux were grown in 3 mL of TSB with chloramphenicol for 16 h. Three biological replicate cultures were inoculated to an OD_600_ of 0.01 into a 125 mL flask containing either 25 ml TSB with glucose, TSB with no glucose, or TSB with no glucose buffered at pH 5.5 with or without palmitic acid as indicated and grown at 37°C with 220 rpm. At hourly intervals, OD_600_ was measured using a spectrophotometer, and samples were withdrawn from each flask for quadruplicate luminescence readings (relative light units, RLU) in a 96 well microtiter plate, using a Cytation 5 multimode plate reader (Agilent Instruments). For each reading, 200 μL of bacterial culture was supplemented with 20 μL of decanal to maximize luminescence. The corrected RLU was calculated as the mean of these measurements minus the RLU for each strain carrying empty pGYlux cultured under identical conditions.

### Data analyses

GraphPad Prism (version 10.4.1) was used to create all graphs and perform statistical analyses in this study. In all experiments, unless otherwise stated, triplicate cultures were used, and data were reported on graphs as means ± standard errors (SEM). One-way analysis of variance (ANOVA) with multiple comparisons, or two-way ANOVA with multiple comparisons was used to test statistical significance depending on the nature of the experiment. Significance was defined as stated in the figure legends.

## Acknowledgements

This work was supported through a Grant to M.J.M. from the Natural Sciences and Engineering Research Council (NSERC) of Canada. T.B.T. was the recipient of Dr. RGE Murray Graduate Scholarship award.

